# Type 1 diabetes mellitus-associated genetic variants contribute to overlapping immune regulatory networks

**DOI:** 10.1101/325225

**Authors:** Denis M. Nyaga, Mark H. Vickers, Craig Jefferies, Jo K. Perry, Justin M. O’Sullivan

## Abstract

Type 1 diabetes (T1D) is a chronic metabolic disorder characterised by the autoimmune destruction of insulin-producing pancreatic islet beta cells in genetically predisposed individuals. Genome-wide association studies (GWAS) have identified over 60 risk loci across the human genome, marked by single nucleotide polymorphisms (SNPs), which confer genetic predisposition to T1D. There is increasing evidence that disease-associated SNPs can alter gene expression through spatial interactions that involve distal loci, in a tissue-and development-specific manner. Here, we used three-dimensional (3D) genome organization data to identify genes that physically co-localized with DNA regions that contained T1D-associated SNPs in the nucleus. Analysis of these SNP-gene pairs using the Genotype-Tissue Expression database identified a subset of SNPs that significantly affected gene expression. We identified 298 spatially regulated genes including *HLA-DRB1, LAT, MICA, BTN3A2, CTLA4, CD226, NOTCH1, TRIM26, CLEC2B, TYK2*, and *FLRT3*, which exhibit tissue-specific effects in multiple tissues. We observed that the T1D-associated variants interconnect through networks that form part of the immune regulatory pathways, including immune-cell activation, cytokine signalling, and programmed cell death protein-1 (PD-1). These pathways have been implicated in the pancreatic beta-cell inflammation and destruction as observed in T1D. Our results demonstrate that T1D-associated variants contribute to adaptive immune signalling, and immune-cell proliferation and activation through tissue and cell-type specific regulatory networks.

**Author Summary:** Although genome-wide association studies have identified risk regions across the human genome that predispose individuals to the development of type 1 diabetes (T1D), the mechanisms through which these regions contribute to disease is unclear. Here, we used population-based genetic data from genome-wide association studies (GWAS) to understand how the three-dimensional (3D) organization of the DNA contributes to the differential expression of genes involved in immune system dysregulation as observed in T1D. We identified interconnected regulatory networks that affect immune pathways (adaptive immune signalling and immune-cell proliferation and activation) in a tissue and cell-type specific manner. Some of these pathways are implicated in the pancreatic beta-cell destruction. However, we observed other regulatory changes in tissues that are not typically considered to be central to the pathology of T1D, which represents a novel insight into the disease. Collectively, our data represent a novel resource for the hypothesis-driven development of diagnostic, prognostic and therapeutic interventions in T1D.

## Introduction

Type 1 diabetes (T1D) is a chronic immune-mediated disease characterised by the progressive loss of insulin-secreting pancreatic beta-cells, and the incidence is slowly rising worldwide [1]. Well-powered genome-wide association studies (GWAS) have identified more than 60 susceptibility loci to T1D which are marked by single-nucleotide polymorphisms (SNPs) [2]. To date, however, the functional roles of most T1D-associated genetic variants is yet to be determined. Notably, 90 percent of these genetic variants fall outside coding regions [3], and therefore, their biological role in the pathogenesis of diseases is not clear. However, there is growing evidence supporting a putative role for these non-coding variants in the regulation of gene expression, as the majority of SNPs fall within regulatory loci such as enhancer regions [4–6].

Classically, GWAS-associated SNPs which fall outside the coding regions of genes have been assumed to affect the most ‘biologically relevant’ or closest genes [7]. A fundamental problem with this assumption is that many intergenic SNPs may influence the expression of genes which are quite distal [8,9]. Indeed, our enhanced understanding of chromosome architecture and nuclear organization over recent years has shown that interactions with regulatory regions (*e.g*. insulators and enhancers) regularly bypass the closest genes and are associated with changes in gene transcript levels of genes located large genomic distances away. These interactions may occur within the same, or on different chromosomes [4,10,11].

Most studies on spatial genomics ignore the impact of these distal-regulatory chromosome interactions despite increasing evidence that genetic polymorphisms as identified by GWAS can alter the expression of genes through distal spatial interactions in a tissue- and development-specific manner [12–14]. For example, Fadason *et al.* identified spatially regulated genes (*e.g. IRS1, ADIPOQ, FADS2, PPA2,* and *WFS1*) within tissues and pathways that are recognized as important for type 2 diabetes progression [12]. This was achieved by integrating information on spatial chromatin organization [15], and functional data (*i.e.* expression quantitative trait loci (eQTL)) to assign SNPs to the genes they control. This finding highlights the importance of integrating an understanding of the spatial genomic context into analyses to discover the fundamental mechanisms underlying gene regulation [4,16–18].

We hypothesised that T1D-associated SNPs contribute to disease pathogenesis by deregulating expression of genes involved in immune activation and response in a tissue-specific manner. In the present study, we used the Contextualize Developmental SNPs in 3D (CoDeS3D) algorithm to perform a combined spatial and functional eQTL analysis to assign T1D-associated genetic variants to the genes they regulate. We identified interconnected regulatory networks of spatially associated T1D eQTLs that affect immune pathways (adaptive immune signalling and immune-cell proliferation and activation). We demonstrate that T1D-associated SNPs have effects in tissues that are not classically associated with T1D, such as liver, brain hypothalamus, and adrenal. These findings provide a novel platform for the development of novel diagnostic and therapeutic interventions.

## Results

### T1D-associated variants form a gene regulatory network

We identified 232 cis- and 66 trans-eQTLs, at an FDR of q<0.05 (S1 Table) for 180 SNPs associated with T1D (p ≤ 9.0 × 10^−6^). The functional physical interactions between T1D-associated SNPs and eGenes (*i.e.* the genes whose transcript levels are associated with the identity of the nucleotide at the SNP position) were represented in a circos plot (Fig 1). Notably, we observed a series of trans-eQTLs that connect into and out of the HLA locus (Fig 1B). The observed cis-and trans-eQTL network for T1D associated SNPs is consistent with a functional role for SNPs in modulating gene expression profiles, rather than protein sequences, that predispose an individual to the development of T1D [24]. The identification of 66 trans-eQTLs, which by definition interact with regions >1Mb apart or on different chromosomes, reinforces the importance of not identifying eGenes on the basis of linear proximity in the absence of observable transcriptional effects due to the specific contexts underlying gene expression.

**Figure 1.**
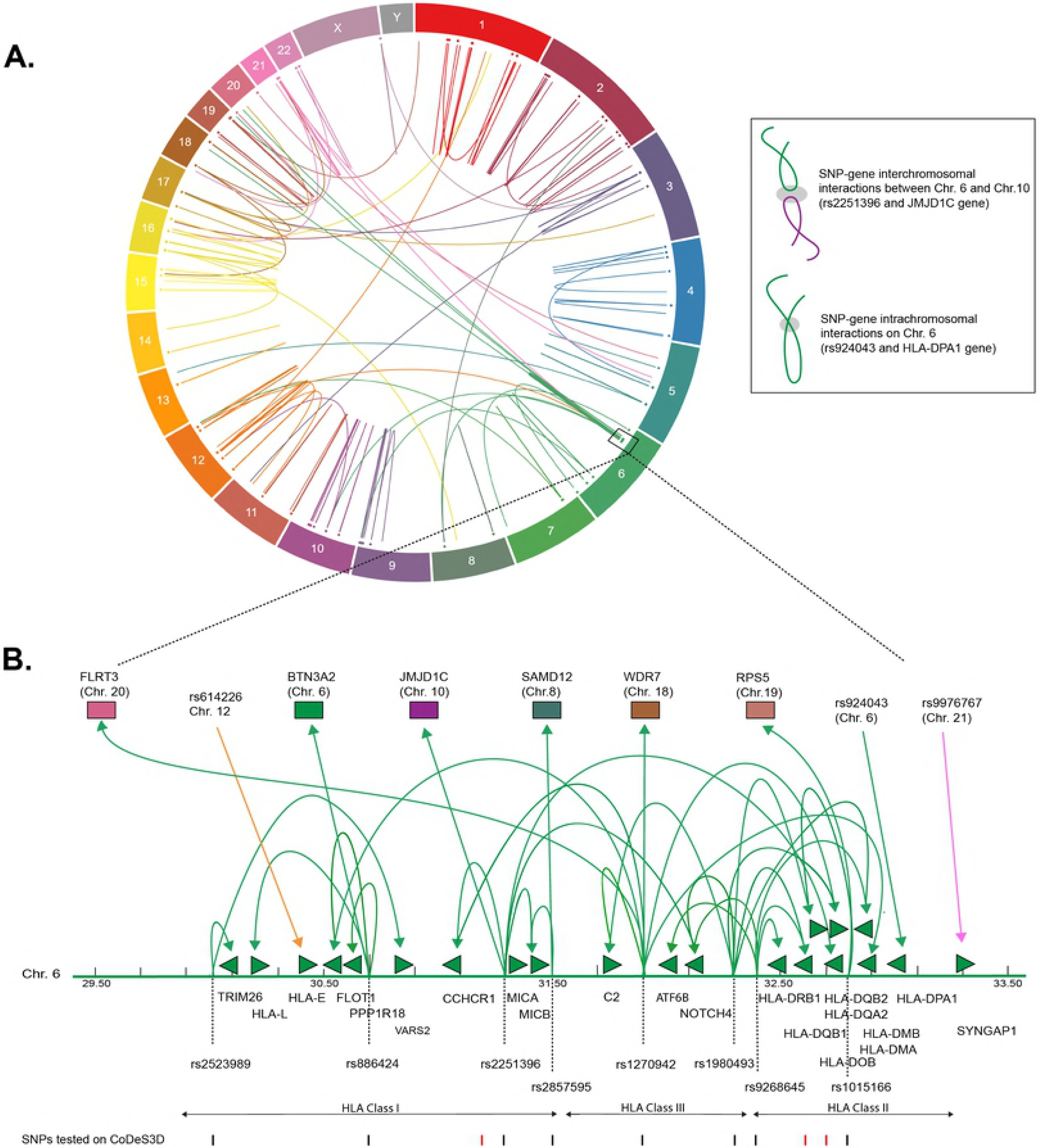
T1D associated SNPs form an integrated gene regulatory network. (A) Circos plot showing interactions between T1D-associated SNPs and significant eQTLs. Data tracks: chromosome labels (outer-most ring); and a scatter plot of relative SNP positions. Link lines represent significant SNP-gene interactions at FDR q<0.05 (S2 Table). The inset illustrates a cis-and trans-eQTL. The grey ellipsoid represents the unknown factors that are responsible for mediating the interaction. (B) T1D associated genetic variants are involved in cis-and trans-eQTLs that enter and emerge from the HLA locus. Significant cis-and trans-eQTLs are annotated by lines with arrows, green arrows heads denoted direction of the regulatory effect. Genes within the HLA locus are annotated as arrows, which indicate transcriptional direction. SNP positions are marked as lines (black - lines indicate significant SNPs, FDR<0.05). eGenes affected by trans-eQTLs are coloured according to chromosome number as in (A).

We used Gene Ontology (GO; http://www.geneontology.org/) and the Reactome Pathway Database to annotate T1D-eGenes (both cis-and trans-eGenes) for biological and functional enrichment. The T1D-eGenes were significantly enriched (FDR, q<0.05) for biological processes and canonical pathways associated with antigen processing and presentation; immune-cell activation (T lymphocytes activity); programmed death signalling; and cytokine signalling (Table 1). Our observations are consistent with the T1D-associated variants modifying the expression of genes that are interconnected (directly or indirectly) through networks that form part of immune system pathways (adaptive immune signalling, and immune-cell proliferation and activation). This agrees with previous observations [5,25], but does not unequivocally prove that variations in these pathways directly contribute to the aetiology of T1D.

**Table 1.**
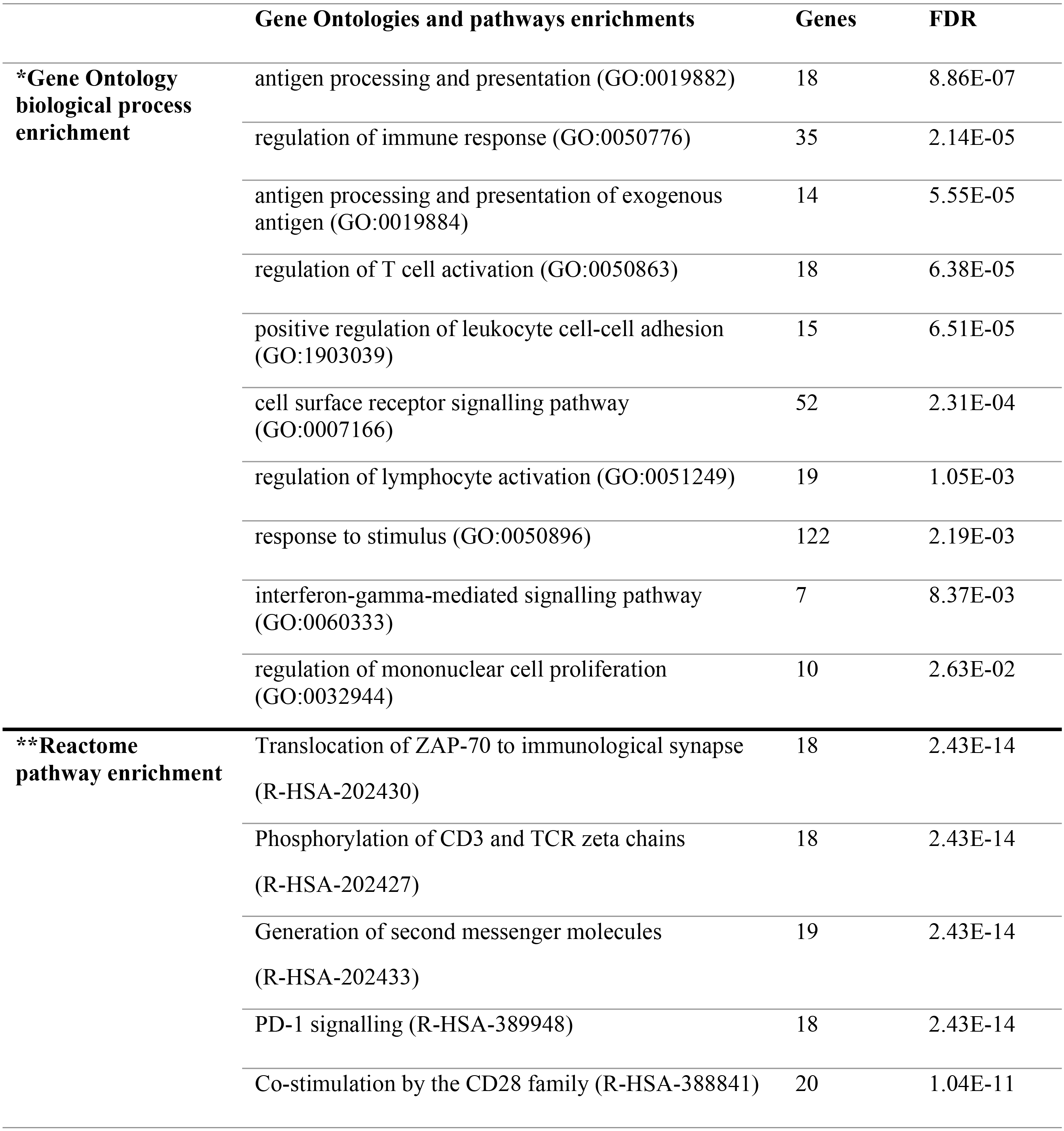
Gene ontologies and pathways enrichment for T1D-eGenes

|  |  |  |
| --- | --- | --- |
| MHC class II antigen presentation (R-HSA-2132295) | 23 | 3.16E-11 |
| Downstream TCR signalling (R-HSA-202424) | 21 | 7.08E-11 |
| TCR signalling (R-HSA-202403) | 22 | 1.65E-10 |
| Interferon gamma signalling (R-HSA-877300) | 27 | 8.26E-10 |
| Interferon signalling (R-HSA-913531) | 29 | 5.58E-07 |

\*Significant gene ontologies (biological process) and \*\*Reactome pathways for T1D-eGenes were derived from http://www.geneontology.org/ and Reactome Pathway Database (version 64, https://reactome.org/) respectively, (FDR $\leq 0.05$ ) [20,22,23,51]. PD-1, programmed cell death protein-1; TCR, T-cell receptor; ZAP-70, Zeta-chain-associated protein kinase 70; MHC, major histocompatibility complex.

### T1D-associated variants across the HLA locus affect transcript levels of genes in cis and trans

There is a possibility that the T1D-associated SNPs located across the HLA locus are associated with the transcript levels of multiple genes or that they combine to regulate the expression of a single strong risk allele. To distinguish between these two possibilities, we analysed the linkage disequilibrium (LD) profile for the HLA based SNPs amongst people with Western European ancestry (CEU). We observed maximal linkage scores of R^2^≤0.6 (Fig 2A). An R^2^>0.8 is generally accepted as indicating robust linkage [26]. Therefore, the linkage we observed between the SNPs we tested was relatively weak (R^2^≤0.6). The inter-eQTL LD we observed is consistent with the majority of the T1D-associated SNPs contributing to the development of disease independently. However, even within the low levels of LD we observed, it is notable that there are two predominant examples of long-distance LD between rs1980493-rs1270942, and rs1270942-rs2647044/rs1980493-rs2647044 (Fig 2A). Long-distance LD has been characterised across the human genome and previously hypothesized to be associated with gene regulation [27].

**Figure 2.**
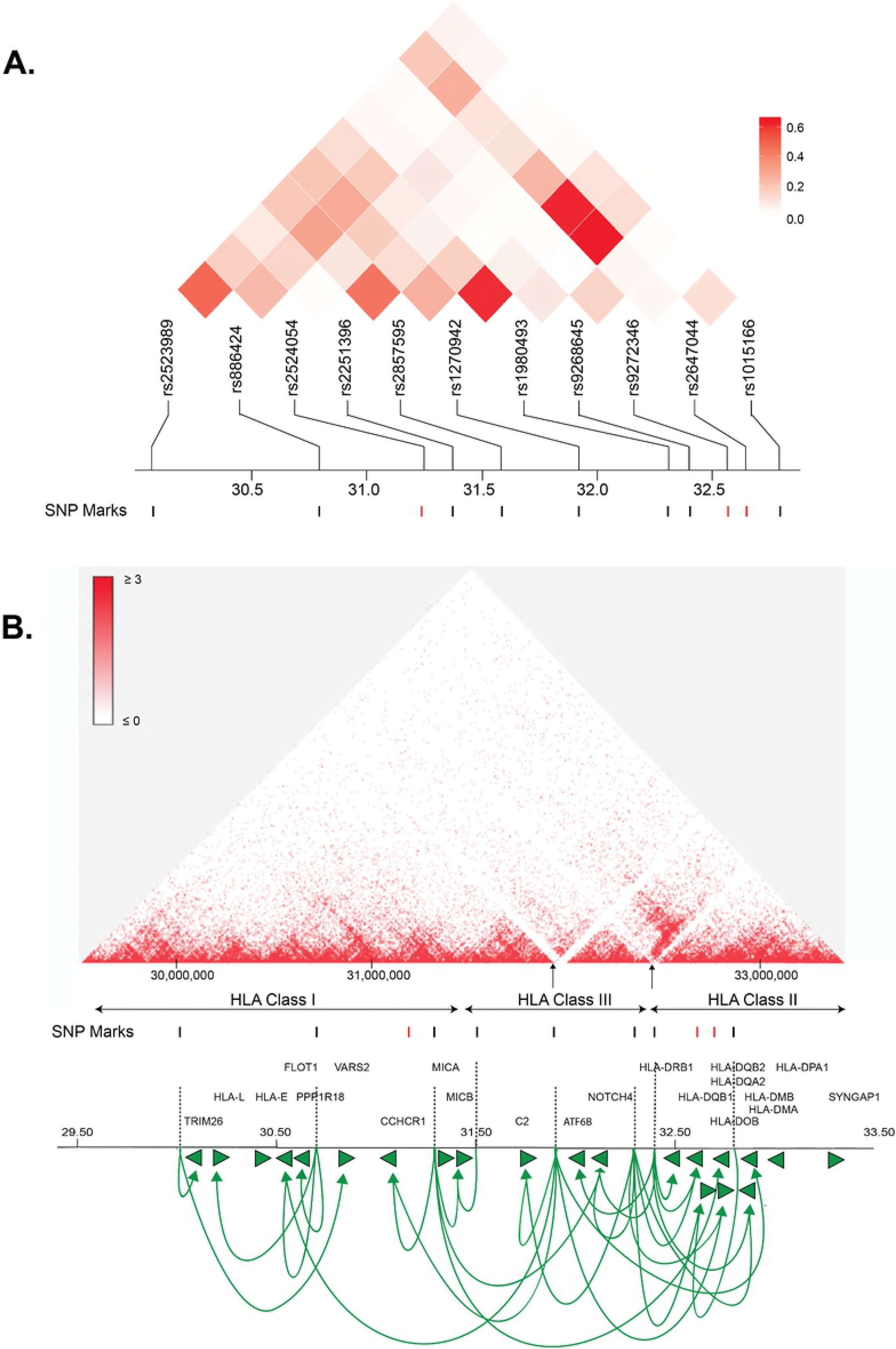
T1D-associated variants along the HLA locus affect gene expression within and outside of the locus. (A) Linkage disequilibrium (LD) plots of T1D-associated SNPs along the HLA locus amongst people with Western European ancestry (CEU). The squares within the heat map represent the LD (R^2^) value between every two variants. The weak LD (R^2^ ≤ 0·6) indicates that SNPs are infrequently co-inherited and contribute to disease development independently. SNPs with significant eQTLs are represented by black marks (FDR ≤ 0·05). (B) A Hi-C heat map of intra-chromosomal contacts across the HLA locus captured in the human lymphoblastoid cell line GM12878 at 10kb resolution [15]. SNPs that show significant and non-significant eQTLs are denoted by black and red marks, respectively (FDR ≤ 0·05). Loops (lines with arrows) represent interactions between SNPs and genes that are associated with differential expression. Green arrows heads denoted direction of the regulatory effect. The two black arrows point to the relative positions of variants rs9268645 and rs1270942 located at or in close proximity to the TAD boundaries. The illustrated gene map is as in Fig 1 (B).

The role of genome structure in gene regulation is widely considered to be represented in the local chromosome structure observed within topologically associating domains (TADs), chromosomal regions which physically interact frequently more than their genomic neighbours [28]. Therefore, we determined how the HLA eQTLs were positioned with respect to TADs and local chromatin structure within human lymphoblastoid cell line GM12878, at 10kb resolution (Fig 2B). The GM12878 lymphoblastoid cell line was used to examine the three-dimensional genome architecture as it has the densest contact map, containing approximately 4.9 billion Hi-C captured contacts [15]. Significant eQTLs involving rs2523989, rs886424, rs2251396, rs2857595, rs1980493, rs1015166 occurred within TADs that were located across the HLA class I, II, and III regions (Fig 2B).

Variants rs9268645 and rs1270942 were located at or in close proximity to the TAD boundaries (Fig 2B). TAD boundary sites have been reported to contain elevated levels of the transcription factor CCCTC-binding factor (CTCF), which is known to act as a chromatin insulator to inhibit transcription and mark transcriptional domain borders [29]. Notably, rs1270942 spatially regulates the expression of genes in both the HLA class I and II regions. Moreover, rs9268645 falls in a boundary region and is an eQTL for HLA-DRB1, whose expression was previously reported to involve regulation by CTCF through long-distance chromatin looping [30].

Three SNPs (rs2524054, rs9272346, and rs2647044) were not found to be involved in eQTLs (Fig 2B). Notably, while rs9272346 and rs2524054 fall within coding regions, rs2647044 is intergenic. This indicates that there remain other as yet undetermined mechanisms through which these genetic variants contribute to T1D disease development. Collectively, our results are consistent with: a) T1D-associated variants located within the HLA locus directly affecting (*i.e.* transcript levels) within and outside of the locus, and b) interactions between functional polymorphisms and gene regulatory elements being associated with inheritance (*i.e*. longdistance LD).

### Expression QTLs contribute to tissue-specific effects in autoimmune T1D

We hypothesised that disease-relevant biological processes fundamentally depended on mRNA levels, with the cross-tissue variability of gene expression providing an important avenue for understanding disease aetiologies. Therefore, we analysed the tissue-specific contributions of T1D-associated eQTLs to tissues. Consistent with our hypothesis, eQTL effects were distributed differently across human tissues (Fig 3A), with tibial artery and lower leg skin (*i.e.* both tissues have been linked to peripheral arterial disease in diabetic patients [31,32]) having the highest proportion; while brain substantia nigra having the lowest proportion of eQTLs (S1 Fig).

**Figure 3.**
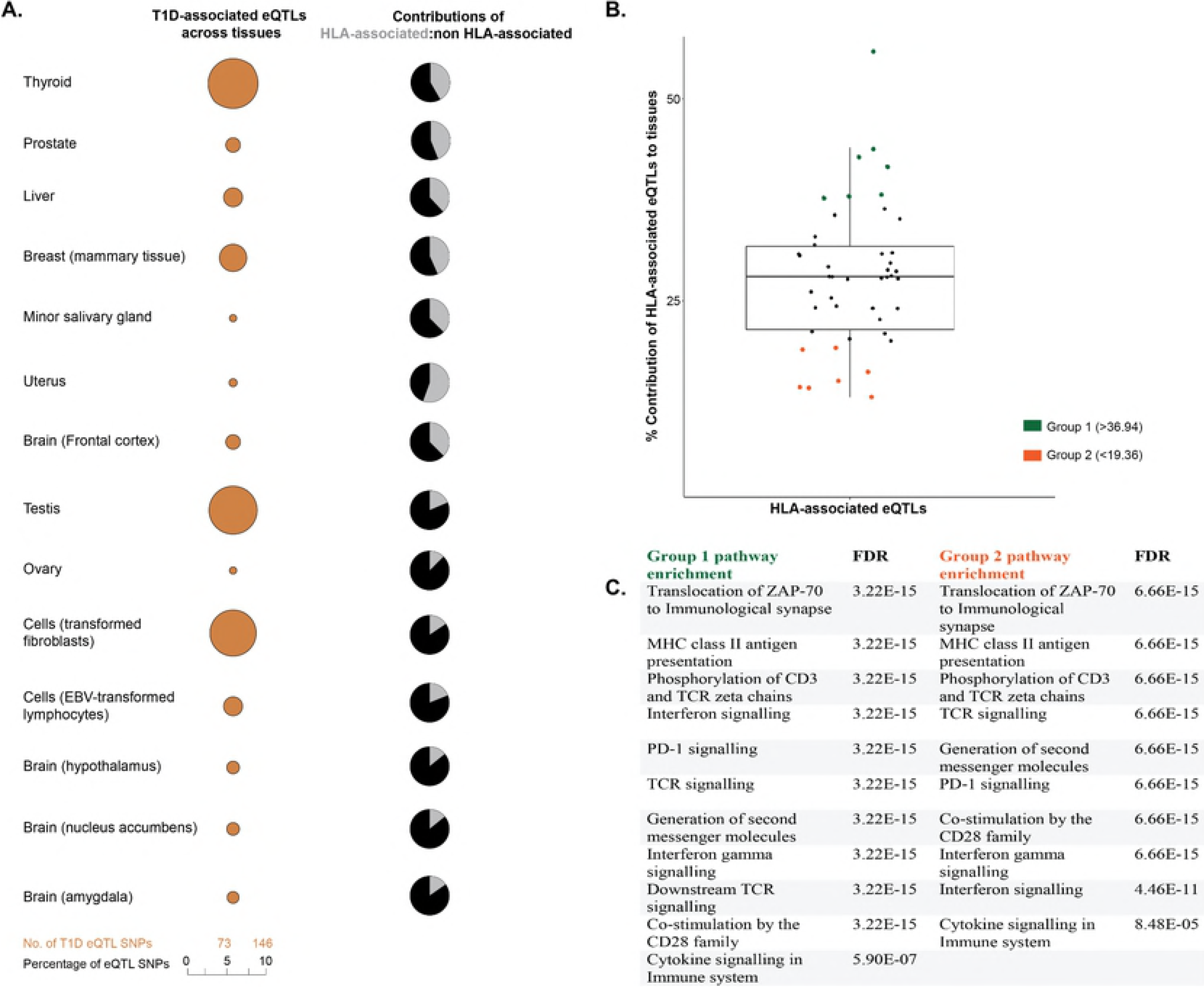
T1D-associated eQTL effects are tissue-specific. (A) T1D-associated eQTLs are differentially distributed across human tissues. The differential distribution is epitomised by the relative proportions of HLA and non-HLA associated eQTLs in different tissues. A complete summary of all GTEx tissues with significant eQTLs (FDR ≤ 0.05) is presented in S1 Fig. (B) The relative contributions of HLA associated T1D eQTLs to tissue specific effects. Relative contribution was calculated (HLA:total eQTLs for a tissue expressed as a percentage). The mean HLA contribution was 28.16 ± 8.79%. (C) eGenes within tissues with high or low HLA contributions (i.e. +/− 1 S.D. from the mean) were enriched for biological pathways associated with immune pathways. Biological pathway enrichment was performed using the Reactome pathways database [22], with significant (FDR ≤ 0.05) for immune response pathways.

We compared the proportions of HLA and non-HLA associated eQTLs across different tissues and identified two tissue groups (group 1 and 2) that were located one SD from the mean (HLA: total eQTL percentage of 28.16 ± 8.79) (Fig 3B). Analysis of the eGenes within these groups, using Reactome pathways database, identified biological enrichment within antigen presentation, programmed cell death signalling, T-cell receptor signalling, and interferon gamma signalling) for both groups of tissues (FDR ≤ 0.05; Fig 3C). The identification of changes in transcript levels involved in immune activation, signalling and response pathways in tissues that are not traditionally associated with T1D pathology is notable. This may indicate that dysregulation of the immune-associated signals within these tissues contributes to the onset or development of T1D. Collectively, we observed cross-tissue concordance for T1D-associated eQTL effects (either HLA or non-HLA associated; Fig 3B), which is consistent with multiple tissues and specific biological pathways impacting on T1D progression [33].

## Discussion

The aetiology of T1D is hypothesised to involve T cell-mediated destruction of the insulin-producing pancreatic islet beta cells, leading to complete insulin loss [1]. We have integrated genetic variation, 3D genome organization and functional analyses (eQTLs) to better understand the downstream effects of SNPs associated with T1D. Our analyses identify associative functional effects of T1D GWAS-associated SNPs in regulating the tissue-specific expression of target genes either through cis- and trans-interactions. Our findings identify overlapping regulatory networks that contribute to adaptive immune signalling, immune-cell proliferation and activation in a tissue-specific manner. We contend that the integration of mixed ‘omic’ datasets into the functional interpretation of T1D-associated SNPs has identified novel pathways and tissue-specific functional genetic loads that represent high priority targets for clinical investigation into the developmental windows and effects that contribute to T1D.

Allelic variations in the antigen presentation mediated by the HLA locus under-pin autoimmune disease [34]. The low levels of LD observed for the eQTLs within the HLA region indicates that these T1D-associated SNPs contribute to the development of disease independently as they are rarely co-inherited. This is consistent with Barcellos *et al*. (2009) who identified the existence of multiple, independent disease susceptibility regions within the HLA locus [35]. However, there are notable examples of multiple susceptibility loci within HLA haplotypes combining to influence T1D risk through co-regulation (Fig 2B). It remains to be determined if the T1D-associated genetic variants within the HLA locus disrupt the coordinated chromatin configuration [34]. However, long-distance regulation involving regulatory loci at TAD boundaries in the HLA locus has been observed previously [30,36]. Future work should use CRISPR-Cas or Degron based strategies [37,38] to empirically confirm the mechanisms by which genetic variations at these loci results in transcriptional changes in order to enable the development of targeted therapeutic or prognostic approaches.

Studies have identified the HLA class II region as a recombination hotspot, resulting in a disrupted LD pattern for a region that is known to exhibit robust linkage [39,40]. Population history (i.*e*. genetic drift), chromosomal recombination activity, and selective pressure could potentially explain the observed long-range LD we observed across the class III - class II HLA regions [40]. We contend that understanding the co-inheritance of the eQTL SNP-egene pairs will provide fundamental insights into the associated protection and susceptibility for particular HLA haplotypes in T1D development across different populations [41]. This will be particular relevant in populations that are undergoing relatively rapid and high levels of genetic admixture.

T1D-associated eQTL mapping studies have focussed on cell-specific effects on immune cell types (*e.g.* [5,42]) based on the assumption that these tissues are critical to understanding the development and pathology of T1D. However, our study strongly implicates dysregulated immune signalling and responses in tissues that are not traditionally considered central to the molecular aetiology of T1D. For example, it remains possible that the high HLA-associated eQTLs functional load that is observed in the uterine tissues reflects the unique immune status of this tissue, and contributes to the development of gestational diabetes mellitus in individuals with a high functional genetic load [43]. Similarly, adipose tissue has recently been hypothesised to play a role in antigen presentation and modulation of T cells [44]. Therefore, identifying the developmental windows and how the functional genetic loads within these non-traditional tissues contribute to dysregulated immune activity in T1D [45,46] will be important in uncovering the trigger(s) for and developmental programme of T1D.

The *BTN3A2* gene product plays an important role in T-cell responses in the adaptive immune response by inhibiting the release of interferon gamma (IFN-γ) from activated T-cells [47]. Aberrant expression of IFN-γ is associated with a pathogenic role in T1D [48]. Therefore, it was significant that we observed that *BTN3A2* transcript levels were linked to rs886424, by a spatial trans-eQTL, in 32 tissues. We also observed that rs3184504 is associated with transregulation of *ARHGAP42* (a member of the Rho GTPase activating proteins) transcript levels. Interestingly, Fairfax *et al.* demonstrated a monocyte-specific trans-association of *ARHGAP24* gene with the DRB1*04, *07 and *09 alleles [49]. Notably, these DRB1 alleles are associated with expression of *-HLA-DRB4,* which encodes the DR53 superantigen in autoimmune disease progression [50]. The *BTN3A2* and *ARHGAP42* eQTLs are examples of trans-regulation involving spatial co-localization of genes and regulatory elements. Such regulations have previously been associated with tissue-specific gene control [12]. Notably, we observed 66 trans-regulatory interactions that tended to occur in a tissue-specific manner. As such, we contend that trans-acting variants are involved in a multifactorial process that enhances discovery of previously unrecognized connections between genes, which is fundamental in elucidating networks and pathways, as well as mechanisms of T1D development.

Immune-regulatory mechanisms operate within tolerance ranges, and if not properly regulated they can promote autoimmune reactions. It is therefore intriguing that T1D-associated genetic variants spatially contribute to distinct overlapping regulatory networks that have the potential to modify the development of the autoimmune phenotype. Notably, the direct and indirect interconnectivity of the T1D-associated eQTLs means that they are capable of influencing immune-response genes expression in a tissue and cell-type specific manner. Untangling these effects requires empirical studies that incorporate expression QTL analyses within precision approaches that illuminate the genetic basis of individual immunological responses. These studies should also refine the mapping strategy for the identification of the regulatory connections by extending the Hi-C data to include additional tissue and developmental stage relevant maps of genomic organization. Integration of these data into clinical studies of T1D will enable individualized mechanistic understanding of treatment response, prognosis and disease development.

## Methods

### Identification of T1D associated SNPs

T1D-associated SNPs were extracted from the GWAS catalog (www.ebi.ac.uk/gwas/; November 3, 2017). Analyses were restricted to SNPs with a discovery p-value (p ≤ 9.0 × 10^−6^), initial sample size >1000, replication sample size >500, and a risk allele frequency of variants of ≥ 0.01. Equally sized sets of control SNPs were randomly selected from the Single Nucleotide Polymorphism database (dbSNP database, Build 151; November 10, 2017) using a Python script.

### Regulatory SNP-gene interactions and eQTLs analyses

We identified genes whose transcript levels depend on the identity of the T1D-associated SNP using the CoDeS3D algorithm (GitHub, https://github.com/alcamerone/codes3d) [12]. Initially, the modular python scripts that comprise CoDeS3D use high-resolution Hi-C data [15] to identify spatial co-localization of two DNA regions, one of which is marked by a SNP. These spatially associating genomic regions are not limited to adjacent regions within the linear DNA sequence [12].

Next, data from the Genotype-Tissue Expression (GTEx) database (version 7, https://www.gtexportal.org/; retrieved on November 19, 2017) [12] is incorporated to address whether spatially associated T1D-SNPs are associated with changes in the transcript levels (eQTLs) of the spatially associated genes. This analysis identifies: a) SNP-gene pairs that are expression QTLs; and b) the tissues in which the eQTL is significant, using the Benjamini-Hochberg correction for multiple testing (FDR, q <0.05) [19]. Although the multiple testing burden of eQTL mapping can bias or misinterpret results, our FDR threshold (q <0.05) has been demonstrated to identify biologically significant associations consistently (*e.g.* [12,14]). Cis-expression QTL SNPs were defined as occurring within loci <1Mb. By contrast, transexpression QTL SNPs were defined as occurring between loci >1Mb apart, or on different chromosomes.

### Gene Ontology (GO) and pathway analysis

We used Gene Ontology (GO; http://www.geneontology.org/) and the Reactome Pathway Database (version 64, https://reactome.org/) to annotate significant eGenes (genes regulated by loci marked by the eQTL SNPs) for biological and functional enrichment [20–23].

## Acknowledgements

The authors would like to thank Keith Godfrey for comments on this manuscript.

## Supporting information

**S1 Table. Summary of the gene regulatory networks for T1D-associated SNPs analysed by CoDeS3D**

*T1D-associated SNPs were identified in the GWAS catalog (download date November 3, 2017; version, v1.0). § Spatial SNP-gene pairs were total number of spatial connections between SNPs and gene regulatory regions, FDR ≤ 0.05. ¶ the total number of eQTL SNP-gene (pairs) interactions, FDR ≤ 0.05 in at least one GTEx tissue. ¶¶ Non-redundant significant eQTL SNP-genes pairs (FDR ≤0.05). † Cis-eQTL SNPs were defined as occurring in loci <1Mb, FDR ≤0.05. †† trans-eQTL SNPs were defined as occurring between loci > 1Mb apart, or on different chromosomes, with a FDR ≤0.05. # eGenes were those expression observed to be affected by an eQTL SNP. ** Random human SNPs were generated using a python library from the Single Nucleotide Polymorphism database (dbSNP build 151; November 10, 2017). A total of 80 randomly generated data sets of 180 SNPs were run through the CoDeS3D algorithm. SEM, standard error of the mean; SNP, single nucleotide polymorphism; eQTL, expression quantitative trait loci; eGene, a gene whose expression is associated with an eQTL.

**S2 Table. Spatial interactions between T1D-associated eQTLs and genes**

Spatial interactions between T1D-associated eQTLs and genes. T1D-associated SNPs were identified in the GWAS catalogue. Non-redundant interactions between SNPs* and genes** are analyses from the computational algorithm (CoDeS3D). Genes expressed at >1.0 Read(s) Per Kilobase of transcript per Million mapped reads (RPKM) in at least one GTEx tissue were included. Cell lines: KBM7-human myeloid leukaemia; K562-chronic myelogenous leukaemia; IMR90-human foetal lung; HUVEC-human umbilical vein endothelial cells; NHEK-primary normal human epidermal keratinocytes.

**S1 Fig. The distribution of disease-associated eGenes across human tissues.**

The graph shows the proportions of eQTL-eGene interactions across tissues (x-axis) in comparison to number of GTEx tissue samples (circles). The enrichment of eGenes in tissues and cell types, such as whole blood, pancreas, thyroid, adipose, and skeletal muscle strongly suggests that T1D-associated variants in regulatory regions have tissue-specific effects since many regulatory regions have also been shown to act in a tissue-specific manner (*e.g.* (57)). The relationship between genetic variation, gene expression, and T1D phenotype does not correspond directly to the number of tissue samples in the GTEx database. Genes expressed at >1.0 Read(s) Per Kilobase of transcript per Million mapped reads (RPKM) were included in this analysis (GTEx version 7, accessed on November 19, 2017).

